# A genome-wide polygenic approach to HIV acquisition uncovers link to inflammatory bowel disease and identifies potential novel genetic variants

**DOI:** 10.1101/145383

**Authors:** Robert A. Power, Christian W. Thorball, Istvan Bartha, John R.B. Perry, Paul J McLaren, Tulio de Oliveira, Jacques Fellay

## Abstract

Polygenic approaches using genome-wide data have been hugely successful in confirming and quantifying the heritability of complex human traits. Here, we highlight their ability to identify potential novel risk variants by looking for variants with pleiotropic effect in genetically overlapping phenotypes.

We used LD Score Regression in a sample of 6,315 HIV+ European individuals and 7,247 controls to test for phenotypes genetically overlapping with susceptibility to HIV-1 infection. Using LD Hub, a web tool that performs LD Score Regression, identified two phenotypes with significant genetic overlap: schizophrenia (rG =0.19, p=0.0007 and ulcerative colitis (rG=0.22, p= 0.0061). We further showed that the genetic overlap between HIV acquisition and schizophrenia is likely driven in part by their shared overlap with cannabis use and sexual behavior. BUMHBOX analyses suggested that these genetic overlaps were driven by genome-wide pleiotropy with HIV acquisition rather than heterogeneity within the HIV acquisition sample. The two diseases identified as genetically overlapping with HIV-1 acquisition have >100 associated variants, and we tested if any of them significantly associated with HIV acquisition. We observed three variants that exceeded our threshold for statistical significance. Two of these were eQTLs in whole blood for genes coding for proteins suspected to be involved in HIV biology: rs1819333 in *CCR6* (p=0.0002) and rs4932178 in *FURIN* (p=0.00033). However, no signal was found for these variants in two smaller African samples totaling 1015 cases and 963 controls, though the mode of acquisition and genetic architecture of these populations differed.

These results highlight the ability to use polygenic methods to gain new insights into complex diseases and identify potential associations with individual variants. Crucially, the leveraging of existing, publically available data makes these methods a cost-effective approach. In this case, our results add to the evidence for the role of risk taking behavior and inflammation of the bowel in HIV acquisition.

**Author Summary:** The biology of what puts certain individuals at greater risk of HIV acquisition is poorly understood. Using several novel polygenic methods, we identify supporting evidence for two important factors leading to acquisition. First, the role of an individual’s genetic predisposition to risk taking behaviours such as number of sexual partners, age at first sexual intercourse drug use, and mental health problems. Second, the role of gut inflammation, in particular a genetic overlap between HIV acquisition with inflammatory bowel disease and the potential role of CCR6 during infection.

## Introduction

HIV-1 affects 34 million individuals worldwide^1^. HIV targets CD4+ T cells for its own replication, leading to a steady decline in immune function, and ultimately manifesting in AIDS if left untreated. While treatment programs have increased life expectancy for those infected^2^, new infections are still high and maintain the epidemic. With only a small minority of sexual encounters leading to infection, understanding what puts an individual at risk of acquiring HIV is crucial to reducing transmission.

Part of the variability in risk of HIV acquisition is known to be due to host genetics^3^. Susceptibility to HIV is associated with a variant within the *CCR5* gene, which encodes the main HIV co-receptor ^4^. The *CCR5-delta32* deletion leads to a smaller protein that is no longer expressed at the cell surface. When present in homozygous form, the *CCR5-delta32* deletion prevents infection by most HIV strains^5^. Multiple genome-wide association studies (GWAS) have been performed to identify novel genetic risk variants associated with HIV-related outcomes. These have consistently identified variants in the MHC as the major host determinant of viral load and disease progression^6−11^, as well as extreme phenotypes such as elite control ^12,13^, long-term non-progression^14^ and rapid progression^15^. By contrast, GWAS of HIV acquisition have not identified reproducible sets of associated genetic variants other than CCR5-delta32. The largest study of HIV acquisition to date, performed by the International Collaboration for the Genomics of HIV (ICGH)^16^, found no genomewide significant single nucleotide polymorphisms (SNPs) contributing to risk of HIV acquisition, suggesting such SNPs do not exist or current studies are underpowered.

However, even when power to detect individual SNPs is lacking, GWAS data can be used to estimate the overall heritability of a phenotype^17^. These so-called polygenic approaches vary in their design, however almost all aim to estimate or predict the extent to which common variants, genome-wide, capture the variation in and across phenotypes. One novel method is linkage disequilibrium score regression (LDSR)^18^. Linkage disequilibrium (LD) is the correlation between SNPs due to coinheritance over generations. GWAS rely on LD to capture a large amount of genetic variation by genotyping common SNPs that correlate with nearby genomic variation. However, SNPs vary in how many other variants they are in LD with, which can be quantified into a SNP’s LD score. This is used by LDSR to address one of the challenges of GWAS: whether systematic inflation in test statistics across SNPs is evidence of confounding or of a polygenic architecture of many causal variants of small effect. In the case of polygenicity, SNPs that are in high LD with many other variants will be more likely to capture causal variants and so high LD scores should correlate with inflated test statistics. In the case of confounding, population structure should affect all SNPs equally regardless of the SNP’s LD score. LDSR calculates the heritability of a trait by estimating the extent to which SNPs with high LD scores have inflated test statistics and so the level of polygenicity. This same approach can be applied to co-heritability between traits, where the test statistics of two traits correlate more for high LD score SNPs^18^, and to biological pathway analysis, where SNPs in each pathway contribute a disproportional amount of heritability^19^. While many polygenic methods for calculating the heritability and co-heritabilities exist (e.g. ^20,21^), LDSR has three key strengths. First, it requires access only to summary statistics from GWAS rather than access to individual-level genotype data. Second, it is not biased by overlapping samples between studies. Third, LDSR can be performed via the webtool LD Hub^22^, which provides users the ability to upload their GWAS summary statistics for easy automated comparison to those from a collection of publically available studies. This makes testing for genetic correlations between phenotypes, and so identifying novel insights into their biology, much more feasible. The strength of such polygenic approaches is that they provide a means of leveraging publically available data to make insights into the genomic architecture of a phenotype without needing to generate new data.

In this study, we apply LDSR to the previous analysis of HIV acquisition by the ICGH^16^ to identify genetic overlap with other publically available GWAS datasets. Of the 24 phenotypes tested, we identified overlapping heritability with two conditions with an inflammatory component, schizophrenia and ulcerative colitis. We followed this attempting to identify pleitropic associations with HIV acquisition by restricting to the known genome-wide significant SNPs for genetically overlapping phenotypes.

## Results

### HIV phenotype analyses

Significant heritability (h2) was observed for HIV acquisition in the ICGH (h2= 0.28, SE=0.047, p=5.40E-09). Cell type annotation was performed for HIV acquisition as per^19^, to identify if specific biological pathways disproportionately contributed to heritability. This analysis found an enrichment for immune cell annotated variants. These explained 2.55 times as much h2 in proportion to number of SNPs (p=0.04).

### Genetic correlation analyses

For genetic correlation analyses, HIV acquisition was tested for genetic overlap with 24 selected phenotypes using LD Hub. Two phenotypes showed significant genetic correlation with acquisition with: schizophrenia (rG= 0.19, p=0.0007) and ulcerative colitis (rG= 0.22, p= 0.0061).

Next, those phenotypes with significant genetic correlation with HIV acquisition were tested to see if they were driven by genome-wide pleiotropy or by overlap with a subset of HIV cases. Here, BUHMBOX was used to perform an analysis of heterogeneity within the cases^25^, making use of their known genome-wide significant SNPs. These SNPs came from the largest and most recent GWAS of these phenotypes, an analysis of 34,241 cases of schizophrenia^35^ and of 86,640 cases of inflammatory bowel disease (which also included ulcerative colitis)^36^. Given the pleiotropy across inflammatory bowel disease (IBD) phenotypes, the suspected role of IBD in HIV acquisition (see discussion), and, on further analyses, the significant genetic overlap between IBD and HIV acquisition (rG=0.19, p=0.01), all IBD SNPs were included.

As missingness was not allowed and discrete genotype calls required, analysis was restricted to Illumina genotyped cases and controls. This left 3,142 cases and 5,264 controls for IBD and 2,233 cases and 4,722 controls for schizophrenia. No significant heterogeneity was found for either analysis, suggesting the genetic overlaps were not driven by heterogeneity in the HIV sample.

### Potential intermediate behaviours between schizophrenia and HIV acquisition

In order to better understand what drove the shared heritability between HIV acquisition and schizophrenia, LDSR was performed on the results of two GWAS of behavioural traits not included in LD Hub. The first was a GWAS of cannabis use^31^, which showed genetic overlap with both HIV acquisition (rG=0.33, p=0.02) and schizophrenia (rG=0.32, p=3.0E-8) but not ulcerative colitis. The second was a GWAS of sexual behaviours which had information on both number of sexual partners (NSP) and age at first sexual intercourse (AFS)^32^. Again, the sexual behaviours showed significant genetic overlap with both HIV acquisition (NSP rG=0.51, p=2.8E-11; AFS rG=-0.42, p=2.9E-11) and schizophrenia (NSP rG=0.27, p=4.8E-15; AFS rG=-0.08, p=0.004), but not ulcerative colitis. Given the known genome-wide significant SNPs for AFS, Mendelian Randomisation was performed to test if AFS predicted HIV acquisition, using the MR Base for two sample MR package (<u>https://mrcieu.github.io/TwoSampleMR/#references</u>) in R^33^ for the 22 SNPs available in both samples. The result was not significant (inverse variance weighted method p=0.21).

### Testing immunological overlap between ulcerative colitis and HIV acquisition

Given that the overlap between ulcerative colitis and HIV acquisition was likely due to a shared immune component, rather than behavioural factors, LDSR was performed using the results of a GWAS of immune cell traits^34^. However, of the 272 immune phenotypes included in this GWAS, no individual phenotype was found to be significantly heritable or share significant genetic overlap with HIV acquisition, most likely due to the small sample size (n=1,629). As a result, the (non-significant) genetic overlaps of both ulcerative colitis and HIV acquisition with each immune phenotype were tested to see if they broadly correlated in the same direction for both diseases. No correlation in the co-heritabilities of ulcerative colitis with immune phenotypes and HIV acquisition with immune phenotypes was observed (correlation 0.099; p=0.21).

### Pleiotropic SNP analyses

To further explore the shared heritability with schizophrenia and ulcerative colitis, their known genome-wide significant SNPs were tested for association with HIV acquisition. After quality control, restricting to the genome-wide significant SNPs from the previously described GWAS of schizophrenia^35^ and IBD^36^ resulted in 147 independent loci available in the ICGH data, leading to a Bonferroni corrected threshold for significance at p=0.00034. Three significant associations were identified. Two schizophrenia SNPs were significantly associated with HIV acquisition. The first, rs4932178 (p=0.0003), is located upstream of the *FURIN* gene and is an eQTL for several genes in blood, most significantly *FURIN* and *FES*^37^. The second SNP, rs662655 (p=0.0002), maps to a gene-poor region of chromosome 1. The top association from the IBD GWAS were two SNPs in complete LD (rs9459882 and rs1819333, p=0.0002). rs9459882 is an intronic variant within *CCR6*, a chemokine receptor involved in the migration of T cells^38^. rs1819333 maps to the 5’UTR region of the Ribonuclease T2 gene *(RNASET2)*, which is involved in RNA binding^39^ and ROS propagation^40^. These SNPs are eQTLs for *RNASET2* and *CCR6* in blood^37^.

### Replication attempt

We next tried to replicate these three associations in two African samples where HIV acquisition phenotypes were available. Given the incomplete overlap in genotyping between samples and different LD structure between European and African populations, all SNPs in the ICGH sample within 500kb of the top association were used in the replication study (N=88 SNPs). Fixed effects meta-analysis found no significant associations after correction for multiple testing in the replication samples. However, only one of the associated SNPs from the discovery sample was available in the replication cohort (rs4932178), and the cohort’s size mean power was low (24%).

## Discussion

Polygenic approaches present a new means to gain insights into the biology of poorly understood phenotypes such as HIV acquisition. Here, we show how LD score approaches can leverage publically available summary statistics from GWAS to identify potential novel SNPs with pleiotropic effect. By identifying phenotypes that both genetically overlap and have a large number of known risk variants, one can narrow down a list of plausible SNPs and so reduce the burden of multiple testing.

Here, we use LD score regression to show a genetic overlap between HIV acquisition and both ulcerative colitis and schizophrenia. Both diseases have known strong associations with variants within the MHC region, which encodes many genes involved in immunity. This region was excluded here due to its complex LD structure. Ulcerative colitis, and IBD more broadly, is an autoimmune disease characterized by an inflammation of the colon. Recently, results from the CAPRISA004 microbicide gel trial demonstrated a link between heightened inflammation and HIV acquisition risk^41^, suggesting that shared inflammatory pathways may influence both ulcerative colitis and HIV risk. Furthermore, a study of α4β7 antibody therapy, a treatment of IBD, showed increased virologic control in monkeys infected with SIV^42^. These previous results, alongside our own findings of a genetic overlap, strongly suggest shared pathogenic mechanisms between IBD and HIV acquisition.

It has also been suggested that schizophrenia has an inflammatory and immune component^43^, though there is debate as to whether the known genetic associations within immune genes could be due in part to their alternate role in neuron communication^44^. A previous study showing the genetic overlap between HIV acquisition and schizophrenia highlighted the role of intermediate behaviours such as risky sexual practices^45^. Our analyses confirm such a hypothesis, showing that both schizophrenia and HIV acquisition genetically overlap significantly with number of sexual partners and age at first sexual intercourse. However, Mendelian randomization did not suggest that AFS directly predicted HIV acquisition. Additionally, schizophrenia is known to associate with behavioural traits such as drug use^46^ that also increase risk of acquiring HIV. Here, HIV acquisition also genetically overlaps with drug use, in the form of cannabis use. Although cannabis use alone is unlikely to increase the risk of HIV infection, these findings likely capture the large shared heritability between substance use phenotypes^31,47^. However, analysis with BUHMBOX showed no evidence of heterogeneity in the genetic overlap between schizophrenia and HIV acquisition. Given only a subset of cases in this sample are likely to have acquired HIV through drug use, this suggests the overlap between these phenotypes was not driven by drug use alone, but a broader behavioural overlap.

The genetic overlaps were used to identify three associated variants potentially involved in HIV acquisition. Of these, two were eQTLs of genes with known or suspected roles in HIV biology. The first association was within the *FURIN* gene, encoding a protein involved in the cleavage and processing of the HIV Tat protein^48,49^. Interestingly, this *FURIN* SNP was associated not just with schizophrenia and HIV acquisition, but also age at first sexual experience, suggesting it may also have a role in the risk taking sexual behaviours. The second association was an eQTL of and intronic variant within *CCR6*, a gene in the same family as *CCR5* within which the well-established delta-32 deletion provides carriers with protection from progression to AIDS. On top of IBD, *CCR6* has been associated with another autoimmune disease, rheumatoid arthritis^50^. The potential role of *CCR6* in HIV acquisition has been a focus of much investigation, with results from animal models suggesting that CCR6+ CD4+ T cells are preferentially infected at the earliest time points during mucosal exposure^51^. Fitting with this, we find that the same allele that increases *CCR6* expression in blood increases risk of HIV acquisition. Independent evidence from hypothesis free GWAS thus provides new evidence for *CCR6* involvement in HIV acquisition, and use as a potential drug or vaccine target.

It is important to note that these associations were not replicated in two independent samples. However, there are several potential explanations for the lack of replication beyond the initial findings being false positives. Due to the ICGH containing the majority of available HIV acquisition samples from European populations, the available replication samples came from Kenya and Malawi. Though the primary modes of acquisition in the ICGH sample were through homosexual contact and injection drug use, the Kenyan and Malawi samples were primarily heterosexually exposed. This means that the routes to acquisition of HIV potentially differed between the discovery and replication samples. Differences also exist in the genetic architecture of the associations, given the different LD patterns in European and African populations and the lack of data for the SNPs of interest in the replication cohort. Lastly, the size of the replication samples, totaling 1,051 cases, was unlikely to have provided sufficient power to detect an effect (24%). As a result, the impact of these associations on HIV acquisition is still in need of further study in appropriately matched samples.

Overall, our results highlight a variety of novel polygenic methods and webtools such as LD Hub, MR Base and BUHMBOX, which can be used to gain new insights into the biology of complex diseases. Of particular interest is their ability to leverage existing, often publically available, data and the possibility of identifying additional genetic risk variants without the need to increase sample size. Specifically, our results provide evidence for the role of risk taking behaviours in HIV acquisition, either resulting from or that simultaneous lead to risky sexual behavior, drug use and mental health problems. Lastly, they support a biological overlap with inflammatory bowel disease, highlighting the potential importance of the gut microbiome, *CCR6* and inflammatory response in HIV acquisition.

## Methods

### Ethics statement

Ethical approval for this study was obtained from institutional review boards at each of the Cohorts, Studies and Centres within the International Collaboration for the Genomics of HIV (ICGH). All subjects provided written informed consent.

### Genome-wide association results from HIV phenotypes

Association results for 9,634,710 SNPs was obtained from a study of the genetics of HIV acquisition performed by the ICGH. The ICGH was established as a platform to combine genome-wide SNP datasets obtained on HIV-1 infected individuals worldwide. Genome-wide SNP data from 6,315 HIV infected individuals of European ancestry was obtained from 25 cohort studies and clinical centers. Genotypes for 7,247 uninfected control individuals matched for ancestry were obtained directly from the participating centres, and from the Illumina genotype control database (www.illumina.com/icontroldb) and the Myocardial Infarction Genetics Consortium (MIGen) (NIH NCBI dbGaP Study Accession: phs000294.v1.p1) ^23^. SNP genotypes were imputed using the 1,000 Genomes Project Phase I (March 2012). All data and quality control and association testing methods are outlined in greater detail in the original paper^16^.

### LD score regression

Summary statistics from the ICGH analyses were uploaded to LD Hub (ldsc.broadinstitute.org) and run against the 24 phenotypes from the original LDSR genetic correlation paper^24^. Analysis was restricted to these phenotypes as they represented a selection of studies screened on ancestry and various QC metrics, as well as being pruned for clusters of highly genetically overlapping phenotypes. For analyses that required LDSR outside of LD Hub, summary statistics of each study were put through the following QC steps where data were available: INFO>0.9, MAF>0.01, removal of small insertions/deletions (indels) and copy number variants (CNVs), removal of SNPs that do not match reference alleles, removal of strand ambiguous SNPs, and removal of SNPs with high missingness, removal of SNPs of extremely large effect. The MHC region (25-35Mb, Chromosome 6) was also removed given its complex LD structure and SNPs of large effect associated with many phenotypes. In studies where the number of individuals for each SNP was not provided, the overall sample size was used. LDSR was performed with the default specifications as in the original publications for estimating heritabilities and genetic correlations^18,19,24^. LD scores for SNPs were downloaded from the LDSR website for HAPMAP3 (github.com/bulik/ldsc).

### Analysis of genetic overlaps

To better understand the cause of any genetic overlaps between HIV acquisition and genetically overlapping phenotypes, we used BUHMBOX to test if genetic correlations were driven by pleiotropy or heterogeneity^25^. BUHMBOX compares LD between known associated SNPs for trait-1 in both cases and controls for trait-2. If SNPs show greater LD in cases than controls it reflects trait-1 risk alleles clustering in a subset of trait-2 cases, rather than being evenly distributed across all trait-2 cases. This suggests that a genetic association between trait-1 and trait-2 is being driven by heterogeneity in trait-2, where a subset of cases overlaps with trait-1, rather than a more general pleoitropy of shared risk alleles across the two traits. In the case of HIV, acquisition can occur in several ways, such as intravenous drug use or sexual transmission, which might overlap with other traits to greater or lesser extents. For genetic correlations observed using LDSR, BUHMBOX analyses were performed using risk alleles from non-HIV phenotype and raw genotypes from ICGH cases and controls. Due to the differing genotyping across studies, analysis was restricted to samples genotyped on Illumina arrays. Genotype files were formatted in PLINK^26^.

### Individual SNP analysis and replication

When phenotypes with significant genetic overlap with HIV acquisition were identified, evidence for pleiotropy on a SNP by SNP basis was tested. Known associated SNPs for genetically overlapping phenotypes were tested for an association with HIV acquisition. P values for genome-wide significant SNPs for genetically overlapping phenotypes were extracted from the ICGH GWAS results. The MHC was again excluded, and only SNPs with an INFO score above 0.9 and a minor allele frequency above 0.05 were included. Significance was assessed after Bonferroni correction.

Replication of significant SNPs was then attempted in two African samples where HIV acquisition data was available. The first was a Malawian cohort of 531 HIV seropositive individuals and 848 highrisk seronegative controls^27^. Briefly, study participants were recruited between 2009 and 2010 in two sexually transmitted diseases clinics in Blantyre and Lilongwe, Malawi. Samples were genotyped on either the Illumina Human1M or 1M-Duo chips. The second cohort was composed of women participating in a well-characterized cohort of female sex workers based in Nairobi, Kenya who were recruited for genetic study^28^. HIV highly exposed persistently seronegative (HESN) women (n=115) were defined as having greater than 3 years of follow-up while remaining active in sex work and HIV negative by serology and PCR. HESN were compared to 484 HIV+ women from the same geographic region. Genotyping was performed using the Affymetrix 5.0 genotyping array and imputed using the 1000 Genomes Project Phase 3 as an extension to previous work^29^. Given the incomplete overlap in genotyping between samples and different LD structure between European and African populations, all SNPs in the ICGH sample within 500kb of the top association were brought forward for replication testing. Fixed effects meta-analysis of the two replication cohorts was performed in METAL^30^. Bonferroni correction for the number of SNPs taken forward for replication was used to determine significance threshold.

## Acknowledgements

The authors have no conflicts of interest. Research supported by a South African MRC Flagship grant (MRC-RFA-UFSP 01 - 2013/UKZN HIVEPI) and Wellcome Trust grant (098051). This research has been conducted using the UK Biobank Resource.

**Figures:**

**Figure 1:**
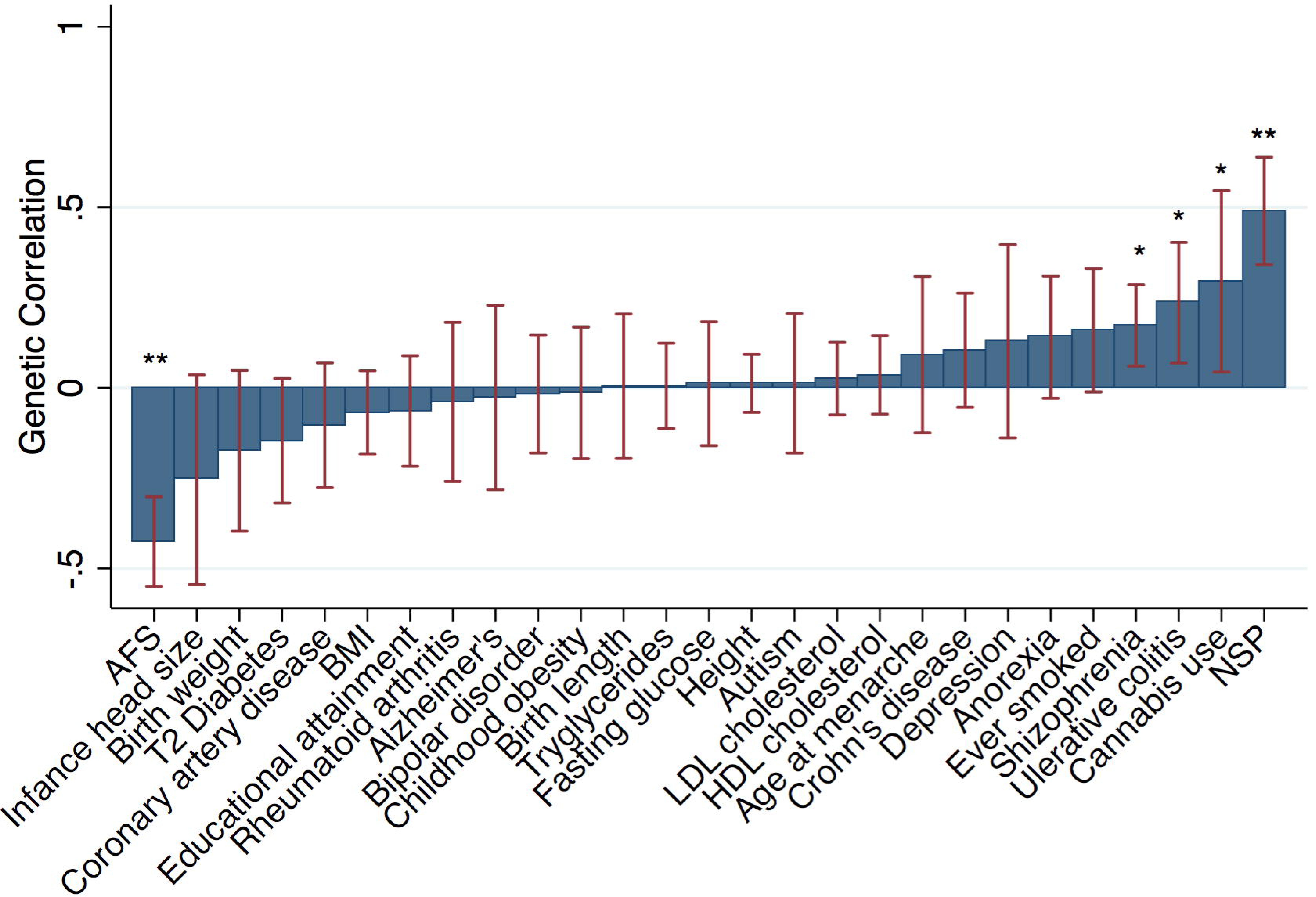
Results of LD score regression analysis of the genetic overlap with HIV acquistion for the selected 24 traits and diseases from the original LDSR paper plus age at first sexual intercourse, number of sexual partners, and cannabis use. * denotes significant genetic correlation at p<0.05; ** denotes significant at p < 1E-5.

**Tables:**

**Table 1:**
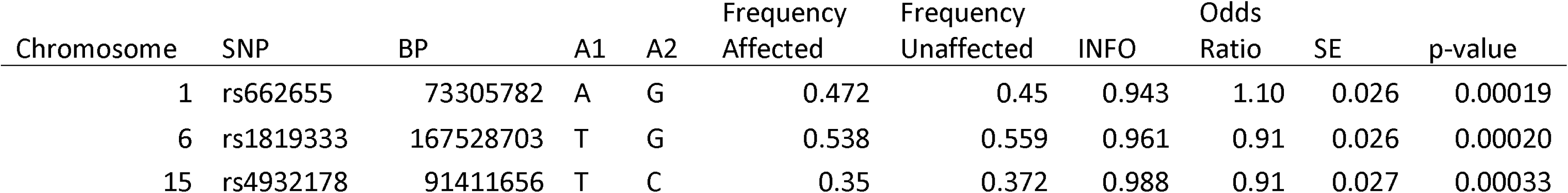
SNPs of pleiotropic effect for HIV acquisition and genetically overlapping phenotypes: schizophrenia and inflammatory bowel disease.

| Chromosome | SNP | BP | A1 | A2 | Frequency Affected | Frequency Unaffected | INFO | Odds Ratio | SE | p-value |
| --- | --- | --- | --- | --- | --- | --- | --- | --- | --- | --- |
| 1 | rs662655 | 73305782 | A | G | 0.472 | 0.45 | 0.943 | 1.10 | 0.026 | 0.00019 |
| 6 | rs1819333 | 167528703 | T | G | 0.538 | 0.559 | 0.961 | 0.91 | 0.026 | 0.00020 |
| 15 | rs4932178 | 91411656 | T | C | 0.35 | 0.372 | 0.988 | 0.91 | 0.027 | 0.00033 |

